# Undoing the ‘nasty: dissecting touch-sensitive stigma movement (thigmonasty) and its loss in self-pollinating monkeyflowers

**DOI:** 10.1101/2024.01.25.577247

**Authors:** Lila Fishman, Mariah McIntosh, Thomas C. Nelson, Kailey Baesen, Findley R. Finseth, Evan Stark-Dykema

## Abstract

Rapid touch-sensitive stigma closure is a novel plant reproductive trait found in hundreds of Lamiales species. The origins, mechanisms, and functions of stigma closure remain poorly understood, but its repeated loss in self-fertilizing taxa and direct tests implicate adaptive roles in animal-mediated cross-pollination. Here, we document several additional losses of stigma closure in monkeyflowers (*Mimulus)*, then use quantitative trait locus (QTL) mapping and gene expression analyses to provide a first glimpse into the genetic and molecular basis of stigma mechanosensing and movement. Variation in stigma closure in hybrids between selfer/non-closer *Mimulus nasutus* and outcrosser/fast-closer *M. guttatus* has a moderately complex genetic basis, with four QTLs together explaining ∼70% of parental divergence.Loss of stigma closure in *M. nasutus* appears genetically independent from other aspects of the floral selfing syndrome and from a parallel loss in M. parishii. Analyses of stylar gene expression in closer *M. guttatus*, *M. nasutus*, and a rare *M. guttatus* non-closer genotype identify functional candidates involved in mechanosensing, turgor regulation, and cell wall remodeling. Together, these analyses reveal a polygenic genetic architecture underlying gain and loss of a novel plant movement, illuminate selfer-outcrosser reproductive divergence, and initiate mechanistic investigations of an unusually visible manifestation of plant intelligence.

## INTRODUCTION

Evolution by natural selection is fundamentally a “tinkerer”, modifying existing genetic material and developmental pathways to build new structures and functions (Jacob 1977). Nonetheless, phenotypic novelty is the spice of life’s diversification (Mayr 1960; Wagner and Lynch 2010; Carscadden et al. 2023), propagating change through re-configured ecological interactions and accelerated radiation within lineages (Miller et al. 2023). Thus, understanding the origins of novel traits remains one of the most fascinating and challenging goals in evolutionary biology. New comparative genomics approaches, combined with functional knowledge from extant organisms, are increasingly revealing the genetic and genomic shifts that contribute to major organismal innovations (Chanderbali et al. 2016; Guijarro-Clarke et al. 2020; Farkas et al. 2022; Clark et al. 2023). A key first step in such analyses is identifying candidate genes and pathways involved in novel trait development through investigating natural losses parallel to the knockout mutations of forward genetics. Although the dismantling of a trait need not follow the same complex path as its construction, the genetics of secondary losses can illuminate both the developmental origins of novelty (Lloyd et al. 2022) and the evolutionary factors important in its maintenance.

In plants, rapid touch-sensitive movement (aka thigmonasty; Braam 2005; Mano and Hasebe 2021) is undoubtedly novel. Plant growth is exquisitely responsive to environmental cues over hours and days and herbivore-damaged cells respond rapidly (Toyota et al. 2018; Kurenda et al. 2019; Farmer et al. 2020; Fotouhi et al. 2022; Matsumura et al. 2022), but acute touch-sensitive mechanical responses are rare. The textbook examples of rapid touch-sensitive movement, trap-closure of carnivorous Venus flytraps (Hedrich and Neher 2018; Vries and Vries 2020) and leaflet-folding of sensitive plant *Mimosa pudica,* are triggered by animals as prey and predators, respectively. This is not surprising, as it is primarily in interactions with moving animals where plant reaction speed matters. Less widely known are thigmonastic movements of floral parts upon contact by animal pollinators (Braam 2005); these include moderately rapid movements of stamens and corolla (Henning et al. 2018; Dai et al. 2021; Li et al. 2022; Tagawa et al. 2022) and rapid (as little as 2 seconds) closure of stigma lobes in many members of the Lamiales (Newcombe 1922). Stigma thigmonasty is widespread, evolutionarily labile (Friedman et al. 2017), and linked to reproductive fitness (Fetscher and Kohn 1999; Krishna et al. 2023), making it an appealing system for understanding the evolutionary genetic of floral mechanosensing and motion. However, despite nearly 150 years of study of touch-sensitive stigma movement (Darwin 1877; Todd 1879; Miyoshi 1891; Burck 1902; Lloyd 1911; Newcombe 1922, 1924), we still know almost nothing about its genetic mechanisms. Revealing the genetic architecture of variation in stigma closure is a key first step toward dissecting its functional basis and investigating its evolutionary origins.

Rapid touch-activated stigma closure is found in Martyniaceae (*Martynia, Proboscidea*; Todd 1879), Linderniaceae (e.g., *Torenia*; Miyoshi 1891), and Mazaceae (*Mazus;* Jin et al. 2015), as well as most members of the Bignoniaceae (e.g. *Incarvillea*; Newcombe 1922, 1924; Ai et al. 2013) and Phrymaceae (*Mimulus*; Newcombe 1922; Friedman et al. 2017) in the Lamiales. This suggests that important genetic and developmental components of the novel trait evolved in a common ancestor. However, taxa with rapid touch-sensitive stigma closure are also interdigitated with families without bilobed stigmas (e.g., Orobanchaceae and Lamiaceae), families with non-closing bilobed stigmas (e.g., Pedaliaceae) and close congeners with slow post-pollination closure of stigma lobes without touch-sensitivity (Newcombe 1924). Furthermore, stigmas with rapid touch-sensitive closure generally also exhibit permanent closure over the hours or days after successful pollination; the latter behavior, which may create a suitable micro-environment for pollen germination or decrease heterospecific pollen-clogging is also found in numerous bi-lobed and trilobed taxa without rapid closure (Waser and Fugate 1986; Webb and Lloyd 1986). This diversity of stigma closure phenotypes suggests that the rapid mechanosensitive movement seen in the most “irritable” species may be built upon ancestral reproductive signaling systems (e.g., changes in stigma and style turgor induced by pollen tube growth) found across angiosperms. Thus, mechanistic investigation of rapid touch-sensitive stigma closure may also provide a window into pollen-style signaling more generally.

Here, we investigate the genetic basis of rapid touch-sensitive stigma closure (TSSC) in monkeyflowers (*Mimulus*; Phyrmaceae) by capitalizing on its loss in self-pollinating taxa. In most of the > 200 species of monkeyflowers, the two lobes of the receptive stigmatic surface fold together within a few seconds of pressure, regardless of pollen deposition. The lobes re-open slowly (∼10-40 minutes) post-touch if un-pollinated, but generally remain permanently closed after successful pollination (Newcombe 1922; Meinke 1992; Fetscher and Kohn 1999; Friedman et al. 2017). Stigma closure speed and completeness varies quantitatively across latitudinal gradients associated with seasonality in the widespread yellow monkeyflower *M. guttatus* (section *Simiolus*, 2n = 28), and TSSC been essentially lost in several selfing species within the *M. guttatus* species complex (Friedman et al. 2017). Parallel patterns were reported in *Mimulus* section *Paradanthus* (2n = 32), with a strong correlation between seedset via autogamous (no pollinator) selfing and stigma-closure time across species (Meinke 1992). This abundant variation, and its association with mating system, suggests active maintenance and fine-tuning by natural selection.

The main adaptive explanations for rapid touch-sensitive stigma closure may also explain its repeated loss with shifts to routine selfing (Webb and Lloyd 1986). First, in animal-pollinated flowers with approach herkogamy (i.e., the stigmatic surface extending beyond the anthers), rapid stigma closure limits pollen deposition to initial contact, precluding costly inbreeding depression due to within- or among-flower self-pollination (Newcombe 1922). Alternatively, within-visit stigma closure may reduce pollen loss due to physical interference with outgoing pollinators, thus conferring male (pollen export) rather than female (seed quality) benefits (Webb and Lloyd 1986). The only direct test of these alternatives used artificial probing of single flowers of the hummingbird-pollinated shrub *Mimulus aurantiacus;* it provided convincing evidence of pollen export costs, but not increased selfing, in flowers with stigmas experimentally prevented from closing (Fetscher and Kohn 1999). However, given that rapid touch-sensitive closure is maintained across hundreds of Lamiales taxa with diverse pollinators, male fitness benefits are unlikely to be the sole and universal explanation. Given an adaptive significance specific to outcross pollination, repeated losses of rapid touch-sensitive closure in selfers may reflect either mutational degradation under relaxed selection or active selection for insensitivity and/or non-closing in selfers. The latter (adaptive) explanation for loss may be particularly plausible in highly specialized selfers such as *M. nasutus* (Fishman et al. 2002), where mature anthers and receptive stigma touch within closed flower buds and stigma closure might interfere with efficient autogamous self-pollination. Addressing how and why closure has been repeatedly lost, as well as probing its origins and molecular mechanisms, requires understanding of the genetic and genomic components of the loss.

We first document the loss of stigma sensitivity and/or closure in three additional highly selfing monkeyflowers, then map its genetic basis in F_2_ hybrids of selfer *M. nasutus* and bee-pollinated *M. guttatus* (Fig. 1), and characterize shifts in stigma/style gene expression associated with loss of closure. The genetics of floral traits and hybrid incompatibilities has been extensively investigated in the *M. guttatus* complex, providing a solid comparative framework for understanding the genetic architecture of our focal trait as well as any confounding factors. Strong segmental synteny between genomes from the *M. guttatus* (2n = 28) and *M. cardinalis* (2n = 16) species complexes (Fishman et al. 2014) also allows direct comparison of candidate QTL regions from this study and a parallel analysis in the latter group (Chen et al., unpubl. MS). Finally, to generate a portfolio of loci potentially involved in rapid touch-sensitive stigma closure both genome-wide and as candidates within QTLs, we characterize patterns of differential gene expression in stylar tissue in the rapidly closing *M. guttatus* parental line relative to noncloser *M. nasutus* and a rare insensitive/non-closing line (IM709) from the same *M. guttatus* population. Together, these analyses reveal the genetic architecture of a novel floral trait through its secondary loss, characterize divergent gene expression associated with TSSC loss and other components of mating system divergence, and open a path toward understanding the molecular mechanisms of both.

**Figure 1.**
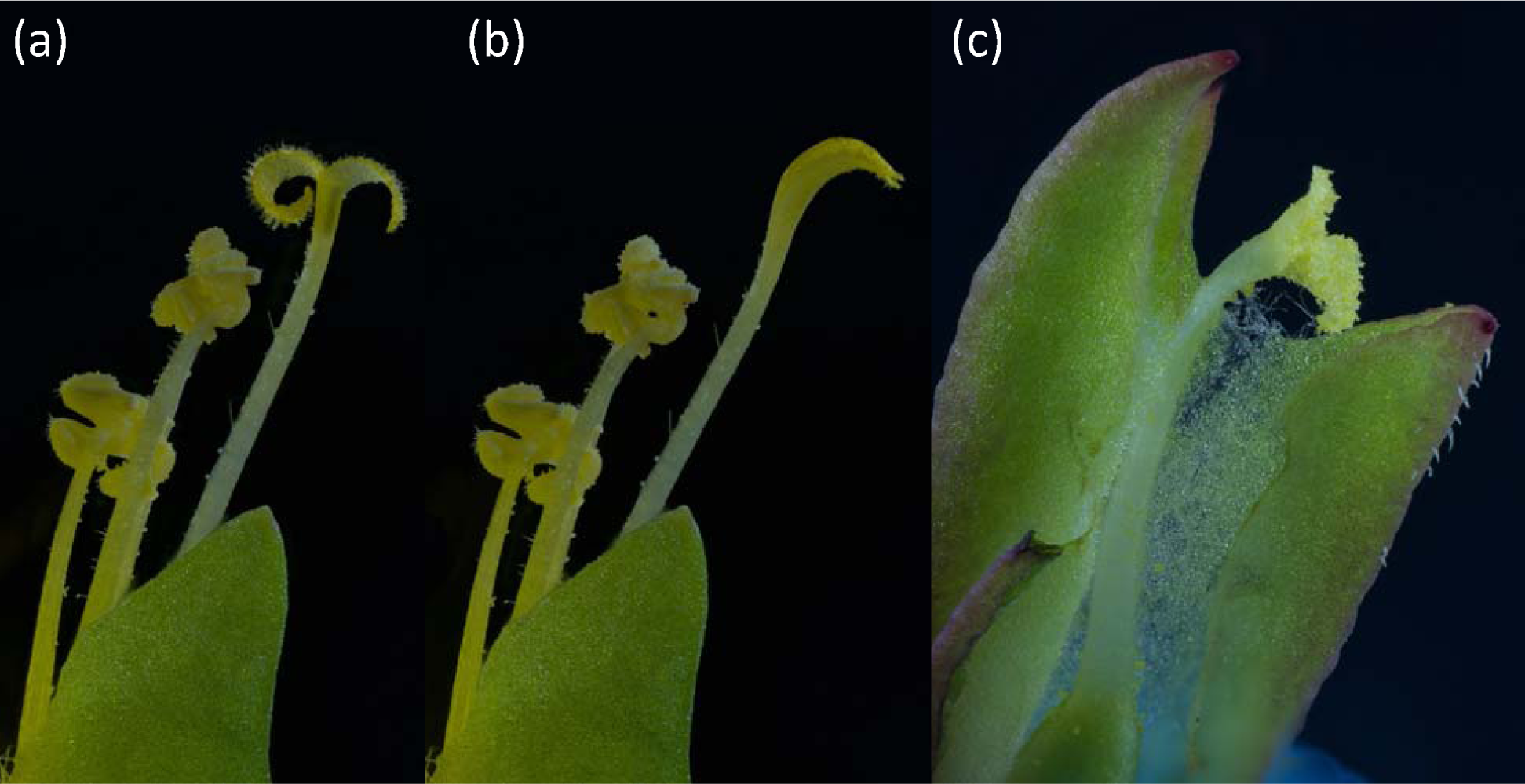
Styles of parental lines of *Mimulus guttatus* (IM767; a, b) and *M. nasutus* (SF: c) used for genetic mapping of the loss of touch-sensitive stigma closure (TSSC) in the latter. (a) Receptive IM767 style, with stamens (corolla removed post-anthesis). (b) Same style ∼5 seconds later after being touched on inner surface of stigma lobes with a glass rod, showing TSSC. (c) *M. nasutus* style with stigma coated in self-pollen. The corolla, anthers (which were in contact with stigma), and half of calyx have been removed. The lower calyx lobe (green at lower right) and stigma lobes are about the same size in both flowers. Photo credit: Timothy Wheeler.

## METHODS

### Documentation of additional losses of stigma closure in selfing monkeyflowers

Monkeyflowers of the genus *Mimulus* (Phrymaceae) exhibit tremendous diversity across Western North America and are a model system for understanding life history, floral, and edaphic adaptation, as well as speciation. Recent taxonomic treatments have split *Mimulus* into >8 genera, renaming the taxa studied here as *Erythranthe.* For continuity and clarity (Lowry et al. 2019), we continue to refer to the focal lines by their *Mimulus* species names. Previous work characterized dramatic reductions in stigma closure speed and completeness in widespread autogamous selfer *M. nasutus* Greene and Sierran selfer *M. laciniatus* A. Gray, as well as substantial variation across annual and perennial ecotypes of outcrosser *M. guttatus* DC (Friedman et al. 2017). To test this pattern across additional selfing yellow monkeyflowers, we grew up inbred lines of *M. micranthus* A. Heller (EBR10) from the Coast Range of California (Puzey and Vallejo-Marín 2014) and *M. hallii* Greene (NRM) from the Rocky Mountains of Colorado (Ivey et al. 2023). These taxa resemble *M. nasutus* in floral reduction, but phylogeographic analyses (Puzey and Vallejo-Marín 2014) suggest that they represent evolutionarily independent derivations of the selfing syndrome. We also characterized *M. parishii,* a small flowered selfer in the *M. cardinalis* complex (Sotola et al. 2023).

Using fast-closing IM767 *M. guttatus* and CE10 *M. cardinalis* lines as controls, we tested the new *M. guttatus* complex selfers and *M. parishii* under standard monkeyflower growth conditions (20/15° day/night temperature cycle, daily bottom-watering) in a Percival PGC40 growth chamber at the University of Montana ECOR Plant Growth Facility. Experimental flowers were emasculated in the bud and the inner surface of the stigma lobes firmly touched once with a rubber pencil eraser to test for a touch-sensitive closure response (see below for more detail).

With the goal of identifying rare *M. guttatus* nonclosers, we separately screened a set (n = 83) of inbred lines derived from the diverse Iron Mountain (IM) annual *M. guttatus* population (Troth et al. 2018) under more variable greenhouse conditions. We used the same phenotyping protocol, but a coarser 3-point scoring system (0 = no closure, 1= slow, 2 = fast). We scored stigma closure on multiple plants and/or flowers per line and calculated a mean score for each line.

### Genetic mapping of *M. nasutus* closure-loss in hybrids with rapid-closing *M. guttatus*

#### Plant materials and phenotyping

We used inbred lines of *M. nasutus* (SF5, Sherars Falls, Oregon) (Fishman et al. 2001, 2002; Brandvain et al. 2014) and annual highly outcrossing *M. guttatus* (IM767, Iron Mountain, Oregon) (Willis 1999; Puzey et al. 2017; Troth et al. 2018) as the parents. We generated an F_1_ hybrid (SF as seed-parent, with emasculation in bud) and self-pollinated a single F_1_ to generate the F_2_ seeds. F_2_ hybrids and parental controls were sown on wet sand in Parafilm-sealed petri dishes, stratified (4°C) for 1 week, and then germinated at ∼25°C under 16hr day/8hr night light regime. Seedlings were transplanted into 2” pots filled with Sunshine #1 soilless potting mix and grown under summer-mimicking conditions (16hr/day of supplemental lighting, ∼27/10°C day/night temperature cycle) in a University of Montana greenhouse in Spring 2018.

On the day the first flower on each plant opened, we recorded the date (flowering time) and measured floral traits (corolla width, corolla length, stigma-anther separation). Prior to the other floral measurements, a single tester touched each stigma head-on with a pencil eraser and scored stigma closure on a 4-point scale (0 = no closure = SF-like, 3 = fast closure = IM767-like, 1 and 2 = slower and faster intermediates, respectively) based on preliminary tests of the parental lines and F_1_ hybrids. For a subset of plants (n = 274 F_2_s), we also collected all four anthers from a later flower into lactophenol aniline blue dye and measured pollen number and viability using a standard protocol (Sweigart et al. 2006).

#### Genotyping

Genomic DNA was extracted from F_2_ hybrids (N = 576) and parental controls using a 96-well CTAB-chloroform extraction protocol and diluted to ∼ 5 ng/ml for library preparation. Following the BestRAD library preparation protocol (Ali et al. 2016), we used a monkeyflower-optimized double-digest restriction-site associated DNA sequencing method (dx.doi.org/10.17504/protocols.io.6awhafe) to generate genome-wide sequence clusters (tags) in each sample, as described in (Kolis et al. 2022). After serial digestion with *BfaI* then *PstI*, half plates of DNA from F_2_ individuals were ligated to biotinylated adaptors with 48 unique in-line barcodes and pooled. Using NEBNext Ultra II kits for Illumina (New England BioLabs, Ipswich, MA), each pool was indexed with a unique NEBNext i7 adapter and an i5 adapter containing a degenerate barcode (for removal of PCR duplicates) and PCR amplified with 12 cycles. The amplified dual-indexed libraries were size-selected to 200-700bp and sequenced (150-bp paired-end reads) in a partial lane of an Illumina HiSeq4000 sequencer at the University of Oregon Genomics Core Facility. After sequencing, samples were demultiplexed using a custom Python script (dx.doi.org/10.17504/protocols.io.bjnbkman), trimmed using Trimmomatic (Bolger et al. 2014), mapped to the *M. guttatus* IM62 v2.0 reference using BWA MEM, and indexed using SAMtools (Li et al. 2009). We followed GATK best practices to create a single VCF with all informative variants, filtered to sites with < 20% missing data and exactly two alleles using vcftools (Danecek et al. 2011), and then thinned to a single informative site per kilobasepair (kb).

We generated linkage maps using Lep-MAP3 (Rastas 2017). First, we removed non-informative sites using the ParentCall2 module. We then filtered out loci deviating from Hardy-Weinberg Equilibrium at *P* < 1×10^-9^; this threshold was empirically chosen to remove rare clusters of “bad” SNPs (generally entire tags with high excess heterozygosity due to cross-mapping of reads among sites). We used the *SeparateChromosomes2* module to assign markers to linkage groups (LodLimit = 65; fixed recombination fraction (θ) = 0.03) and manually combined linkage groups belonging to the same chromosome on the reference assembly. Next, we performed iterative ordering using the *OrderMarkers2* module (Kosambi mapping function; 6 iterations/per linkage group), manually pruned end markers that drastically increased map lengths, and re-ordered all pruned linkage groups; the order with the highest likelihood for each linkage group was chosen. The resulting linkage map and Lepmap3-smoothed (Rastas 2017) genotype matrix consisted of 3284 markers. We manually re-oriented and numbered linkage groups to match previous genetic and physical maps, then pruned to a subset of markers with unique positions (n = 806) for QTL mapping.

We mapped QTLs for floral traits using composite interval mapping (window size =10 cM, background markers = 10) implemented in WindowsQTLCart, following a previous study of mating system traits in *Mimulus* (Fishman et al. 2015). LOD thresholds for QTL detection were set for each trait separately with 1000 permutations. Because both floral differentiation and hybrid fertility have been extensively characterized in SF *M. nasutus* x IM62 *M. guttatus* hybrids (e.g., Fishman et al. 2002; Sweigart et al. 2006) and are included here as possible correlates of stigma closure, we used a genome-wide *P* = 0.05 threshold for stigma closure QTLs and an additional *P* = 0.10 threshold for other traits.

### Analyses of gene expression in styles and stigmas of closers and nonclosers

To compare stylar expression profiles, we harvested whole styles from replicate plants of the three focal lines raised in the growth chambers. To prevent pollen contamination, we removed the corolla and anthers of sample flowers prior to pollen release, following established emasculation for each species. For *M. nasutus*, the underside of the tubular calyx and corolla were slit with forceps and epipetalous stamens carefully dissected out prior to the corolla bud showing any color beyond the calyx. For IM767 and IM709 *M. guttatus*, the corollas of closed flower buds were pulled off one day prior to opening, removing the not-yet-mature epipetalous stamens. After 24 hours, stigmas were inspected for stray pollen grains and damage under a Leica dissecting scope. Clean and intact styles (but no green ovary tissue) were flash frozen with liquid N_2_ in 1.5 ml tubes (n = 8-12 styles per tube, n = 3 tubes per genotype) and homogenized by hand-grinding with disposable pestles in-tube. RNA was extracted using Zymo Plant RNA kits (Zymo Research, Irvine CA), DNased, and quantified using an Agilent TapeStation (Agilent, Santa Clara, CA). Non-directional, polyA-enriched RNASeq libraries were prepared using a PolyA selection NEBNext Ultra II RNA kit (New England BioLabs, Ipswich MA) and sequenced on the Illumina NovaSeq S4 sequencing platform (2×150 bp reads) at Admera Health Corporation (South Plainfield, NJ).

For expression analyses, the de-multiplexed fastq data files were trimmed and filtered using Trimmomatic v. 0.35 (Bolger et al. 2014), then aligned with STAR 2.5.0a (Dobin et al. 2013) to a custom pseudo-reference constructed to contain both the IM62 *M. guttatus* V2 genome and SF *M. nasutus* SNP variation for all v2 genes, as previously described (Kerwin and Sweigart 2020; Finseth et al. 2022). Alignments were converted to bams, indexed, and filtered (removed alignments with quality <20, unmapped reads, and non-primary alignments) with SAMtools (Danecek et al. 2011). Duplicate reads were removed with Picard’s ‘MarkDuplicates’ command and reads spanning splice junctions split using GATK ‘SplitNCigarReads’ command in GATK version 4.0.11.0 (VanderAuwera and O’Connor 2020). Read counts for each sample and pseudo-reference gene were generated with Htseq 2.0 using the htseq-count command (Putri et al. 2022). At this point, we excluded one *M. nasutus* samples due to obvious contamination with IM reads (i.e., an excess of reads mapping to the *M. guttatus* half of the pseudo-reference). Read counts were then summed across the *M. guttatus* and SF *M. nasutus* versions of each v2 gene for each remaining sample.

For identification of highly style-expressed genes and analyses of differential expression (DE) in DEBrowser v1.28.0 (Kucukural et al. 2019), we first filtered the full dataset to genes with >5 counts per million in at least 2 samples. To account for differences in library size among samples (range = 6.58-20.73 million reads), we normalized readcounts using MRN (median ratio normalization). This batch-correction resulted in strong separation of the three genotypes as clusters along major principal component axes that explained 38% (*M. guttatus* vs. *M. nasutus*) and 29% of the variance (line), respectively. To rank stylar expression of genes within each line, we calculated the mean of the batch-corrected readcounts for each gene and standardized these values by the length of the longest transcript to account for gene size variation (custom scripts: https://github.com/FishmanLab-UM/Stigma_RNA), and then examined the most highly expressed genes (top 50, or ∼0.3%). For comparing these genes to a previously published style (minus stigma) proteome (Aagaard et al. 2013), we translated their v1 genome annotations to v2 names, joined with the 15,510-gene expression set, and compared standardized mean readcounts to peptide abundances using correlation analysis in JMP 17 (SAS Institute, Cary, NC USA).

For the individual non-closer vs. IM767 DE comparisons, we further filtered each gene set to those with >5 counts per million in at least 2 samples. We then used likelihood ratio tests implemented in DESeq2 (Love et al. 2014) for separate comparisons of normalized counts of each noncloser to the IM767 closer, using an FDR-adjusted cut-off of *P* = 0.01 to call genes as significantly DE. We used ShinyGO version 0.77 (Ge et al. 2019) to identify gene ontology terms enriched in the significantly up- and down-regulated gene sets in each of the non-closer vs. IM767 differential expression analyses, using the sets of style-expressed genes as background and FDR-adjusted *P* < 0.05 threshold for enrichment.

## RESULTS

### Stigma closure has been independently lost at least three times in selfing monkeyflowers

Selfers *Mimulus micranthus* (EBR10 line), *M. hallii* (NRM line), and *M. parishii* (PAR line) all exhibited no touch-sensitive rapid stigma closure under controlled growth conditions. Outcrossing control plants (*M. guttatus* IM767 line, *M. cardinalis* CE10 line) grown side by side exhibited their usual rapid (< 5 seconds) closure under these experimental growth conditions. This represents at least three independent and parallel losses, as *M. micranthus* and *M. parishii* are distinct lineages from the *M. nasutus* previously tested, and *M. hallii* is likely another independent derivation of selfing within the *M. guttatus* complex.

We identified several slow- or non-closing lines in the Iron Mountain (IM) *M. guttatus* inbred line set. The most extreme, IM709, was as almost as consistently non-closing as SF *M. nasutus*, while three others (IM413, IM266, z453) were often non-closing but exhibited slow stigma closure in some flowers when re-tested. IM709 is similar in floral and vegetative morphology to fast-closer IM767 and the population mean (Troth et al. 2018); however it carries other rare traits (e.g., ability to flower under 12-hour daylengths) that suggest a history of introgression from *M. nasutus* may account for its lack of stigma closure. The remaining quantitative variation should be amenable to genome-wide association mapping in the full set of inbred IM lines with more tightly controlled environmental conditions and increased line-replication.

### Genetic mapping of *M. nasutus* closure-loss in hybrids with rapid-closing *M. guttatus*

#### Parental differences, F_2_ phenotypic correlations, and genetic mapping context

As expected, the IM767 *M. guttatus* parent had significantly higher pollen number, corolla width, and stigma-anther distance (but not flower length) relative to *M. nasutus,* and flowered ∼3 days later. In the F_2_ population, flowering time was positively correlated (i.e., later = higher values, all *P* < 0.01) with corolla width (*r* = 0.14), stigma-anther separation (*r* = 0.36), and pollen number (*r* = 0.18), but not with stigma closure speed (*P* > 0.05). The highly correlated measures of corolla size (*r* = 0.68) were positively correlated with stigma closure score (*r* = 0.15 and 0.16, *P* <0.0003). Mapped markers on the 14 *M. nasutus* x *M. guttatus* F_2_ linkage groups (total length = 1237.4 cM) spanned 92% of the *M. guttatus* v2 genome assembly (Fig. 2). No informative markers in the first 10Mb of Chr3 accounted for most of this deficit, and the adjacent region of LG3 exhibited excess transmission of *M. guttatus* alleles (57:43 IM:SF vs. 50:50 expectation; *P* < 0.001). Beyond the *inv10* region of recombination suppression the map is freely recombining and generally consistent with the *M. guttatus* v2 genome order.

**Figure 2.**
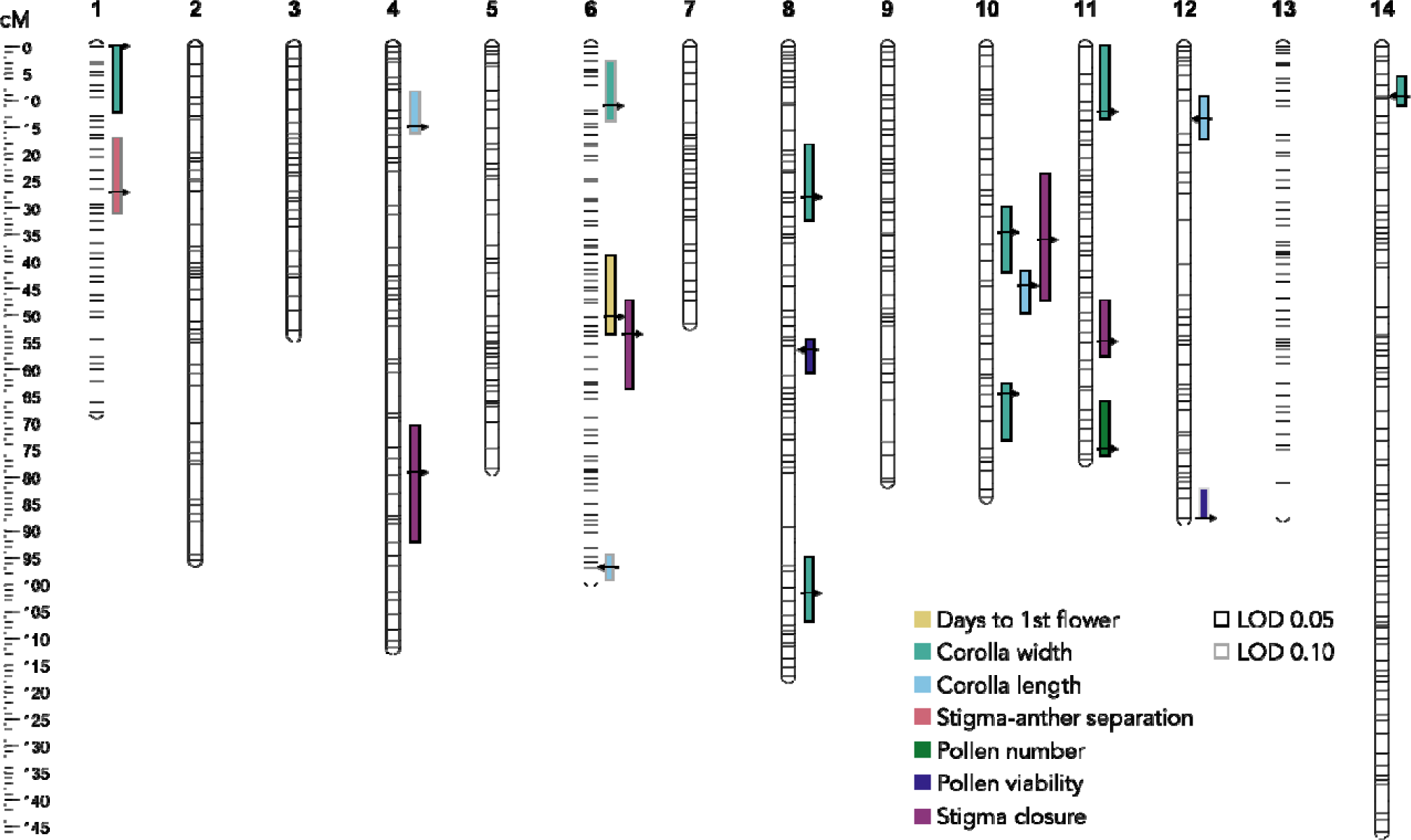
Quantitative trait loci (QTLs) for stigma closure speed and other floral traits on the SF *M. nasutus* x IM767 *M. guttatus* linkage map (14 linkage groups = chromosomes, total length = 1237.4 cM). Stigma closure QTLs were identified at 0.05 LOD threshold only, while other traits were assessed at 0.05 and 0.10 LOD thresholds. Bars show QTL confidence intervals (1.5 LOD drop), with arrows indicating peaks (pointing right if *M. guttatus* alleles increase the trait value, left if the opposite). Further information on QTL effect sizes is in Table 1.

**Table 1.**
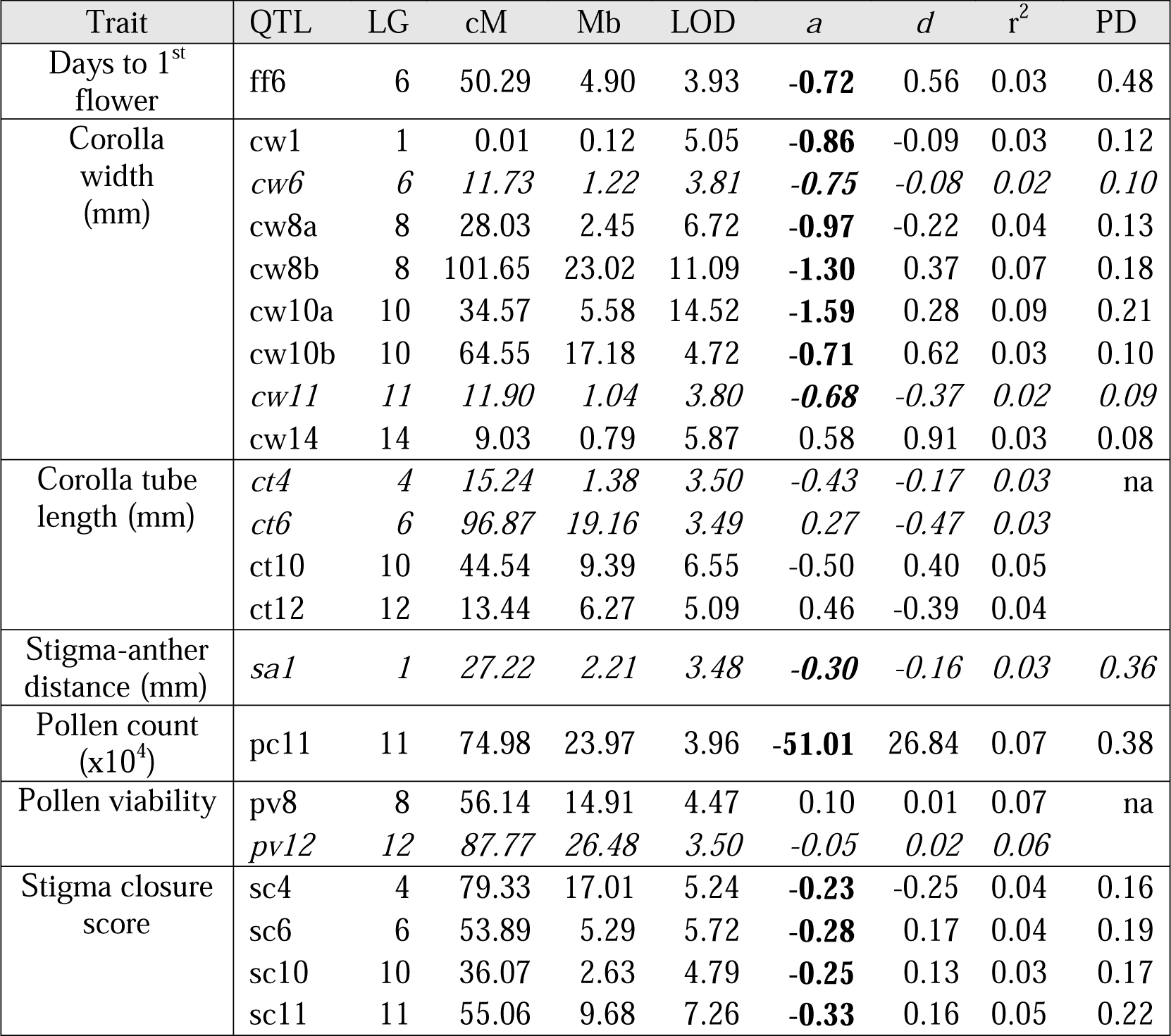
Quantitative trait loci (QTLs) for floral traits, pollen fertility, and touch-sensitive stigma closure in *Mimulus nasutus* x IM767 *M. guttatus* F_2_ hybrids. QTL peaks are named and localized by the linkage group (LG; also the chromosome number), centiMorgan position (cM), and chromosomal position in the *M. guttatus* v2 genome in megabases (Mb). The statistical strength (logarithm of odds ratio; LOD), additive effect (*a*), dominance effect (*d*), proportion of the F_2_ variance explained by (r^2^), and proportion of the mean difference between parental lines explained by 2*a* (PD) estimate the magnitude of QTL effects. QTL information is italicized for floral QTLs identified only at a genome-wide *P* = 0.10 threshold and additive effects are bolded if they move the phenotype in the direction expected from the mean difference between the parental lines (which were not significantly differentiated for corolla length or pollen viability; na = not applicable).

#### QTLs for flowering time, floral dimensions, and fitness traits

Floral divergence between *M. nasutus* and IM *M. guttatus* is due to minor loci, with no individual QTL in this study explaining >9% of the F_2_ phenotypic variance (Fig. 2, Table 1). Corolla width was most polygenic, with eight QTLs together explaining ∼1/3 of the F_2_ variance. In contrast, four corolla length QTLs cancelled each other out, consistent with no flower length difference between *M. nasutus* and the IM767 *M. guttatus* line. We detected only one QTL each for flowering time (LG6) and pollen number (LG1) and only a very marginal QTL for stigma-anther separation (LG1; LOD = 3.49, genome-wide *P* = 0.12). The opposite effects of *M. nasutus* homozygosity at the two pollen viability QTLs (*pv8* significant, *pv12* marginal), along with no fertility difference between the parents, point to a weak recessive-recessive Dobzhansky-Muller incompatibility independent of the polymorphic LG6/LG13 *hms1/hms2* interaction identified in hybrids between *M. nasutus* and the IM62 *M. guttatus* line (Sweigart et al. 2006; Sweigart and Flagel 2015).

#### QTLs for stigma closure

We detected significant QTLs for stigma closure score on LG4, LG6, LG10 (coincident with *inv10* at ∼32-36cM), and LG11(Fig. 2, Table 1). *M. nasutus* alleles decreased stigma reactivity at all QTLs and accounted for more than 70% of the closure difference between the parents. At each significant QTL, one homozygote (*M. guttatus* for *sc4*, *M. naustus* for the others) was significantly different from the other two genotypes by Tukey-Kramer HSD tests. To visualize the distribution of closure phenotypes for multi-locus genotypes, we re-coded the peak genotypes as *M. nasutus* dominant (*sc4*) or recessive (*sc6, sc10, sc11*) to generate a composite score ranging from 0 (all *M. nasutus;* mean = 0.67) to 8 (all *M. guttatus;* mean = 2.24) (Fig. 3) A test for pairwise epistasis among QTL genotypes (ANOVA in JMP, with all four QTLs plus the pairwise interactions) found only weak interaction between *sc6* and *sc11* genotypes (*P* = 0.04).

**Figure 3.**
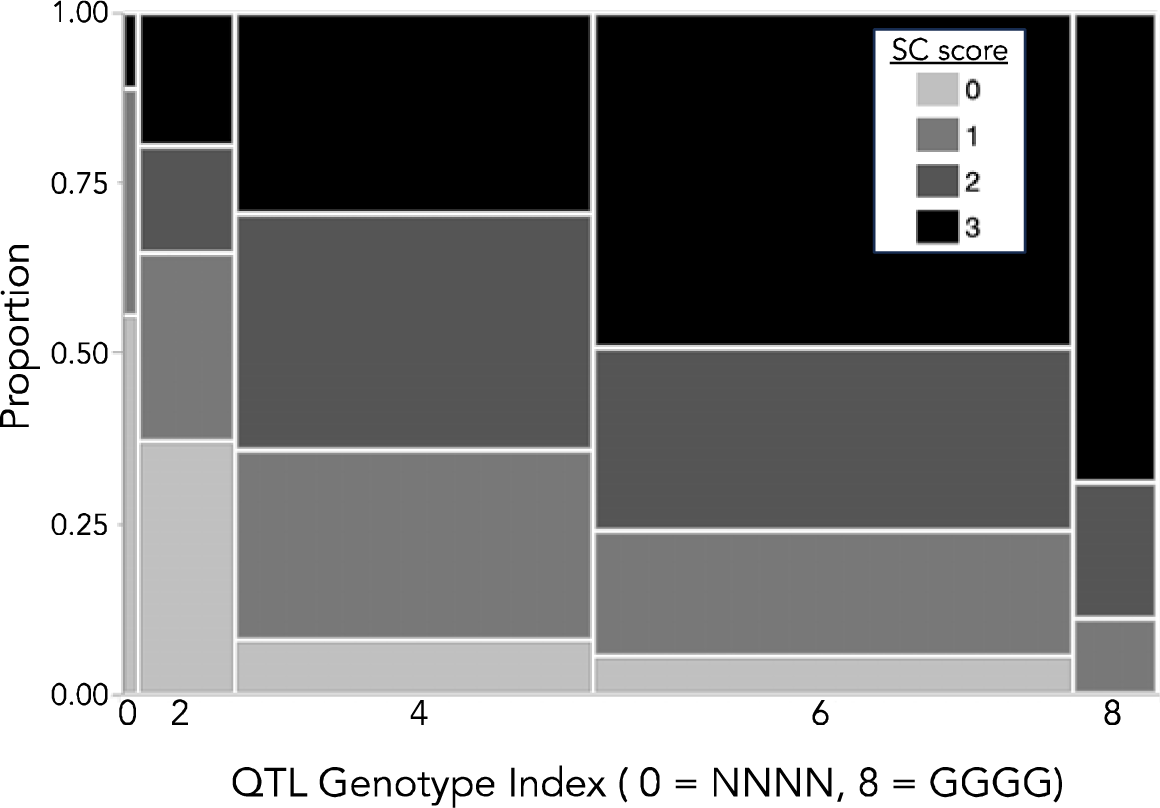
Four quantitative trait loci (QTLs) cumulatively affect stigma closure speed in hybrids between noncloser *M. nasutus* and fast closer IM767 *M. guttatus*. F_2_ hybrid genotypes at QTL positions have been recoded to reflect QTL dominance (0 = *M. nasutus*-like, 2 = *M. guttatus*-like) and summed across the four loci to calculate a QTL Genotype Index. Width and length of each block indicate the sample size of F2s with a given index value and the proportion with each stigma closure (SC) score respectively.

### Highly expressed stylar genes in *Mimulus* closers and nonclosers

We retained 15,510 genes (∼55% of genome-wide total) as style-expressed in at least one of the three genotypes. The top-50 most expressed genes in the three lines were highly overlapping, with 24 of the 86 total genes shared by all three lines, 16 shared by a pair, and additional functional overlap even when the exact genes were not shared (Table S1). The highly expressed set shared by all lines included a Mechanosensitive Channel of Small Conductance-like 10 (MSL10) on Chr 1 (Migut.A00554), three Plasma Intrinsic Protein 1 (PIP1) family aquaporins (Migut.F00699, Migut.F01419, and Migut.B01762), four lipid transfer proteins (LTPs; Migut.A00823, Migut.H01994, Migut.J00043, and Migut.J00046), and three pectin lyases (Migut.F00292, Migut.O00218, Migut.B00929), two unlinked homologues of the ethylene-forming enzyme ACO4 (Migut.H01595 and Migut.M01363) and a MYB21 transcription factor implicated in cell elongation. Eight additional aquaporins (both PIPs and tonoplast intrinsic proteins or TIPs), five additional LTPs, one more pectin lyase, three terpene synthases, and three expansins involved in cell elongation, and other genes with inferred functions in cell wall remodeling and biotic and abiotic stress responses, were found in one or more top-expressed set.

We also cross-referenced the style transcriptome with genes identified as components of the N^15^-labelled stylar proteome in a study focused on *M. guttatus* pollen tube proteins (Aagaard et al. 2013); 94% of the proteome genes (n =2484, including some non-unique) were present in the transcriptome dataset and standardized mean readcounts of the three lines were all positively correlated with the Multidimensional Protein Identification Technology (MuDPIT) normalized peptide abundance in IM62 *M. guttatus* (r^2^ = 0.10-0.19, all *P* < 0.0001, n = 2344) in that study. However, only <50% of the genes in the top-50 set (46/86) were present in that stylar proteome, which excluded stigma tissue (Table S1).

### Differential expression between nonclosing selfing species *M. nasutus* and closer *M. guttatus*

Genome-wide, 4229/14,672 retained genes were differentially expressed, with 2191 genes up-regulated in *M. nasutus* relative to IM767 and 2038 genes down-regulated (Figure 4). The set with elevated expression in *M. nasutus* was significantly enriched for nitrate transporters (12/46), aquaporins (12/28) and xylanase inhibitor/aspartic peptidase (19/127) and glycoside hydrolase (49/239) family genes associated with cell wall remodeling, as well as broad cytochrome P450 and transferase categories. The DE subset with low expression in SF was enriched for several categories associated with cell division, growth, and maintenance (MADS box, Chaperonin/TCP, ribosome biosynthesis, RNA processing), as well as defense against pathogens (NB-ARC) and organellar transcript processing (pentatricopeptide repeat genes; PPRs). Notably, terpene synthase genes (8/27) were nearly 6-fold over-represented; three homologs of terpene synthase (TPS21) that specifically generates stylar volatiles in *Arabidopsis* were in the top-50 genes in IM767 but low expressed in both SF *M. nasutus* and IM709. The highly expressed MSL10 on Chr1, while a strong functional candidate for stigma mechanosensing, was not differentially expressed in this comparison.

**Figure 4.**
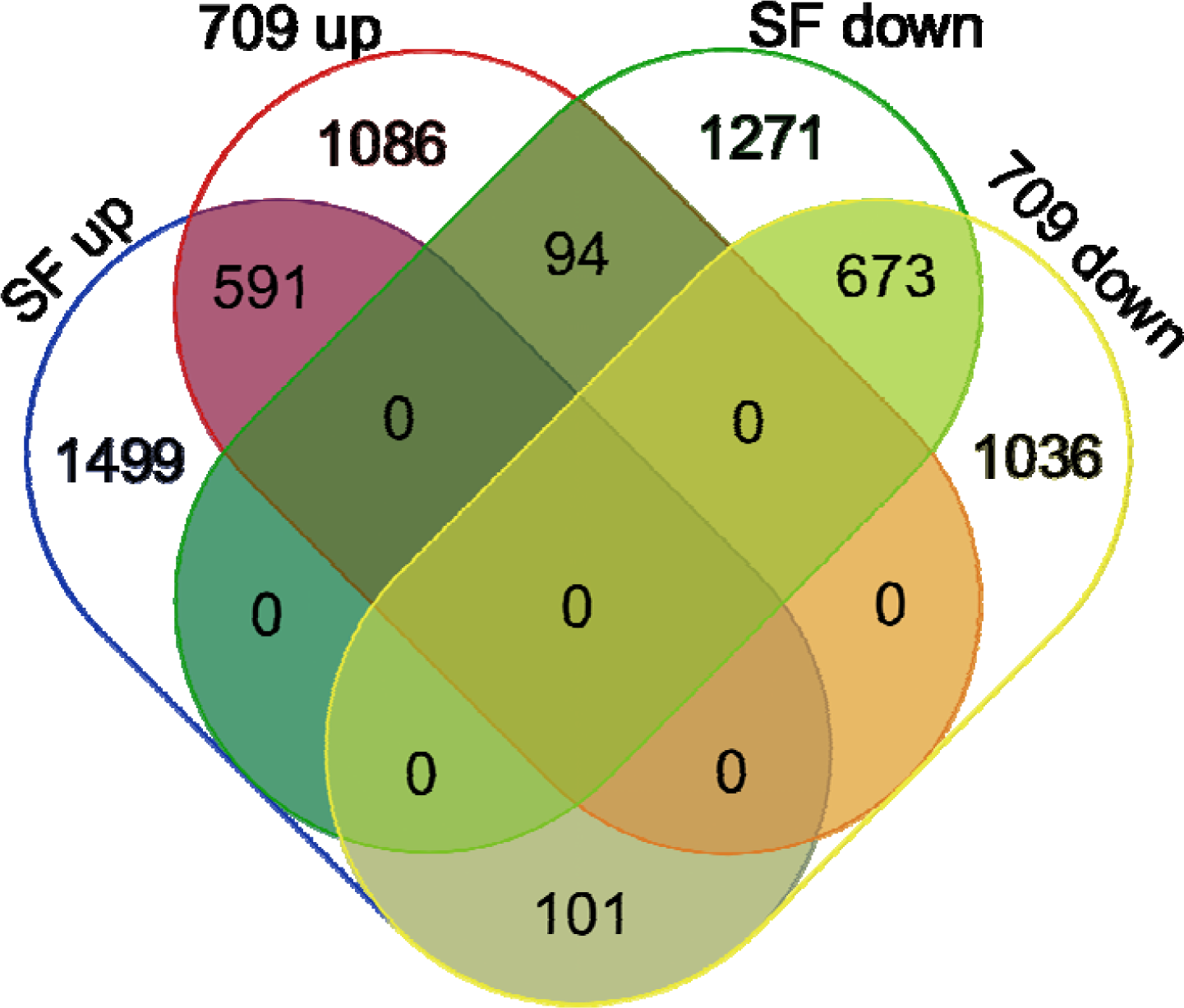
Overlap of differentially-expressed style gene sets in comparisons of IM767 *M. guttatus* (fast stigma closure) with SF *M. nasutus* (no closure) and IM709 *M. guttatus* (no/slow closure). The regions with zeroes (up/down overlaps within a line) are impossible combinations. Venn diagram drawn at https://bioinformatics.psb.ugent.be/webtools/Venn/.

Nearly a third (300/1017) of style-expressed genes in the four stigma closure QTL regions were significantly differentially expressed, leaving many genes in play as dual genetic expression candidates. Under the *sc4* QTL, Migut.D02133 (*sc4*), a tonoplast intrinsic protein (TIP) aquaporin with 8.5-fold higher expression in *M. nasutus,* mirrors the genome-wide aquaporin pattern, and may be involved in the maintenance of turgor. In the *sc6* QTL, tandem UDP glucosyltransferase 72 genes (Migut.F01070 and Migut.F01071) may be candidates for cell wall modifications; in *Arabidopsis*, knockdown of the corresponding gene causes ectopic lignification and thickening of cell walls in the inflorescence stem (Lin et al. 2016). Migut.J00949, a Mechanosensitive Ion Channel 10 (MSL10) family gene is a candidate within the *inv10/sc10* QTL; it is weakly expressed relative to the MSL10 on Chr 1 but shows >2.5-fold higher expression in IM767 vs. *M. nasutus*. Under the *sc11* QTL, Migut.K00910 (a homologue of Arabidopsis Receptor-like Kinase 1, which is involved in aquaporin regulation) is 18-fold higher expressed in *M. nasutus* and may contribute to both loss of closure and the gene-wide shifts in aquaporin gene expression.

### Differential expression between rare noncloser and closer lines of IM *M. guttatus*

In the intra-population comparison between rare noncloser *M. guttatus* line IM709 and rapid closer IM767, somewhat fewer genes (3581/14,678) were differentially expressed (Figure 4). The set with higher expression in IM709 vs. IM767 styles was enriched for nitrate transporters (12/85) and transferase (13/104), tyrosinase (6/11), and AMP-dependent synthetase (11/38) families, as well as cytochrome P450 and PPR genes. The set lower in IM709 vs. IM767 expression set was >5-fold enriched for terpene synthases (7/27), diacylglycerol kinases (7/11), and weakly enriched for Leucine-rich repeat genes (Table 2). The highly style-expressed MSL10 on Chr1 (Migut.A00554) was significantly but weakly differentially expressed in this contrast (FDR-adjusted *P* = 0.003), with 2-fold higher expression in IM767 relative to IM709. More than 40% of the IM709-IM767 DE genes were shared with the larger SF-IM767 DE set, just 13% (195) of those showed DE in opposite directions (Figure 4), and joint DE genes showed highly correlated shifts in expression (*r* = 0.72, P < 0.0001). The striking shared reduction in terpene synthase expression, which includes >20-fold reductions in three terpene synthase 21 (TPS21) homologs from the IM767 top-50, suggests a shared loss likely independent of stigma closure. TPS21 in *Arabidopsis* is a style-expressed producer of sesquiterpene floral volatiles, which are key to pollinator attraction in *M. guttatus* (Haber et al. 2019) and may also contribute to bacterial defense. Relaxed selection on both functions is a plausible secondary consequence of routine autogamous selfing, which makes bees and the microbes they carry inconsequential. Other shared enrichment patterns, particularly the parallel reduced expression of transporter genes and enrichment for pathways involved in cell-wall remodeling in both comparisons, suggests consistent gene-expression modules associated with loss of stigma closure.

**Table 2.**
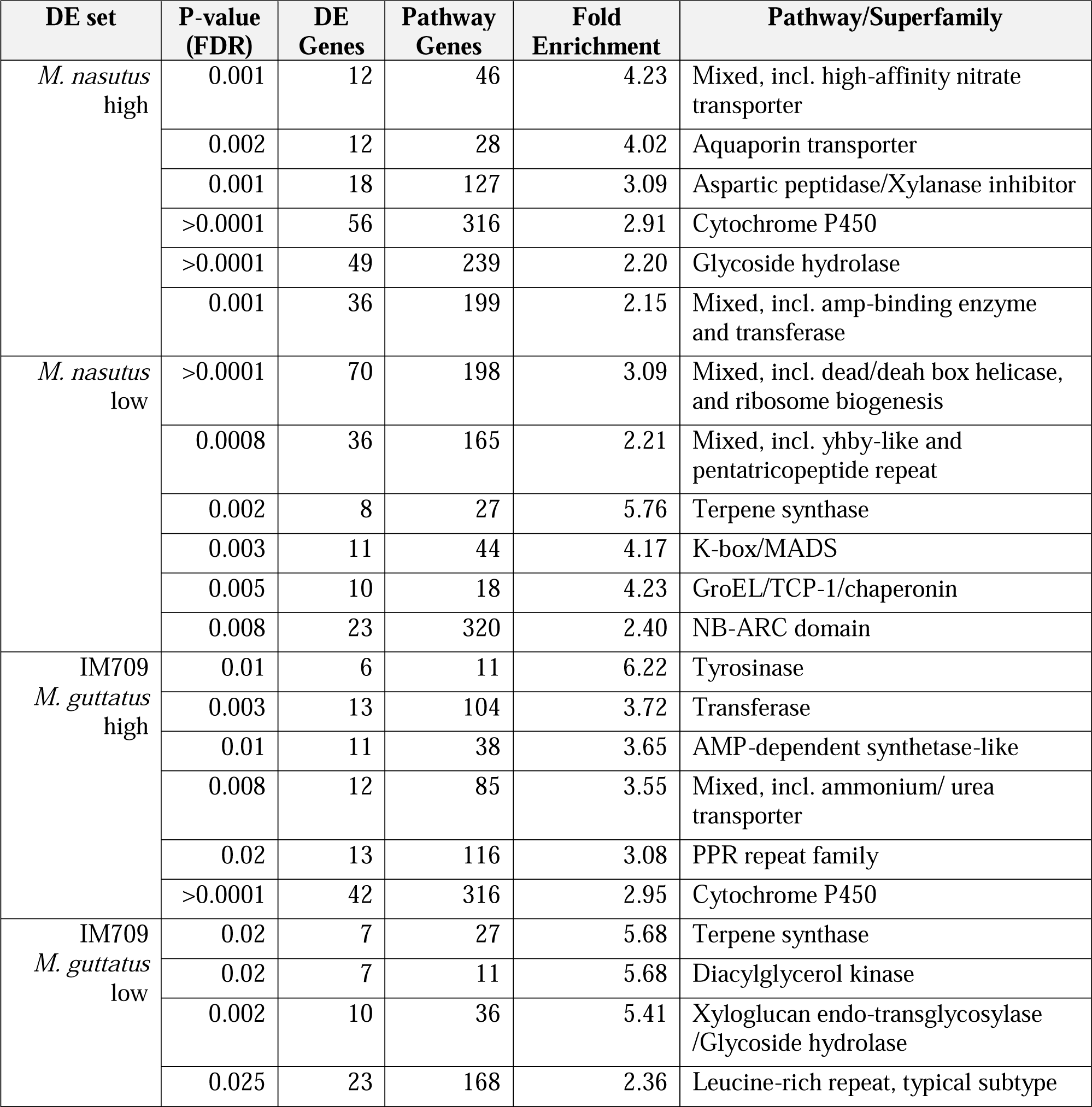
Pathways/gene families enriched for differentially-expressed stylar genes in comparisons of nonclosers SF *M. nasutus* and IM709 *M. guttatus* to fast closer IM767 *M. guttatus.* Enriched pathways were slimmed to a single representative member (usually the most specific) using the tree/clustering outputs in ShinyGO 0.7.7.

| DE set | P-value (FDR) | DE Genes | Pathway Genes | Fold Enrichment | Pathway/Superfamily |
| --- | --- | --- | --- | --- | --- |
| *M. nasutus*<br>high | 0.001 | 12 | 46 | 4.23 | Mixed, incl. high-affinity nitrate transporter |
|  | 0.002 | 12 | 28 | 4.02 | Aquaporin transporter |
|  | 0.001 | 18 | 127 | 3.09 | Aspartic peptidase/Xylanase inhibitor |
|  | >0.0001 | 56 | 316 | 2.91 | Cytochrome P450 |
|  | >0.0001 | 49 | 239 | 2.20 | Glycoside hydrolase |
|  | 0.001 | 36 | 199 | 2.15 | Mixed, incl. amp-binding enzyme and transferase |
| *M. nasutus*<br>low | >0.0001 | 70 | 198 | 3.09 | Mixed, incl. dead/deah box helicase, and ribosome biogenesis |
|  | 0.0008 | 36 | 165 | 2.21 | Mixed, incl. yhby-like and pentatricopeptide repeat |
|  | 0.002 | 8 | 27 | 5.76 | Terpene synthase |
|  | 0.003 | 11 | 44 | 4.17 | K-box/MADS |
|  | 0.005 | 10 | 18 | 4.23 | GroEL/TCP-1/chaperonin |
|  | 0.008 | 23 | 320 | 2.40 | NB-ARC domain |
| IM709<br>*M. guttatus*<br>high | 0.01 | 6 | 11 | 6.22 | Tyrosinase |
|  | 0.003 | 13 | 104 | 3.72 | Transferase |
|  | 0.01 | 11 | 38 | 3.65 | AMP-dependent synthetase-like |
|  | 0.008 | 12 | 85 | 3.55 | Mixed, incl. ammonium/ urea transporter |
|  | 0.02 | 13 | 116 | 3.08 | PPR repeat family |
|  | >0.0001 | 42 | 316 | 2.95 | Cytochrome P450 |
| IM709<br>*M. guttatus*<br>low | 0.02 | 7 | 27 | 5.68 | Terpene synthase |
|  | 0.02 | 7 | 11 | 5.68 | Diacylglycerol kinase |
|  | 0.002 | 10 | 36 | 5.41 | Xyloglucan endo-transglycosylase /Glycoside hydrolase |
|  | 0.025 | 23 | 168 | 2.36 | Leucine-rich repeat, typical subtype |

## DISCUSSION

Rapid touch-sensitive stigma closure (TSSC) is a novel plant reproductive trait found in hundreds of species across the Lamiales, including most members of the Phrymaceae and Bignoniaceae. The origins, mechanisms, and functions of stigma closure remain poorly understood, though its repeated loss in self-fertilizing taxa and a few direct tests implicate adaptive role(s) in animal-mediated cross-pollination. The mapping and gene expression analyses of the loss of touch-sensitive stigma closure in self-fertilizing monkeyflowers provide a first glimpse into the genetic and molecular basis of this fascinating plant mechanosensing and movement trait. QTL mapping provides genetic insight into how loss of stigma sensitivity/closure relates and compares to other components of the floral selfing syndrome, while expression analyses illuminate both its mechanism and self-outcrosser functional divergence more broadly. We found a complex genetic basis (in terms of QTL magnitude, number, and gene content) underlying loss of stigma closure in selfer *M. nasutus* (vs. *M. guttatus*) here, as well as in a parallel analysis of *M. parishii* x *M. cardinalis* hybrids (Chen et al., unpubl. MS). Genome-wide patterns of expression in closer and non-closer styles pointed to candidate genes for the ancestral gain of stigma mechanosensing, secondary losses of stigma closure, and the complex shift to the floral selfing syndrome.

### Loss of touch-sensitive stigma closure – an independent component of the selfing syndrome

The shift from outcross pollination by animals to routine autogamous self-pollination is one of the most common evolutionary transitions in flowering plants (Barrett and Harder 1996). The floral syndrome typical of selfers includes reductions in stigma-anther separation (herkogamy) and temporal separation of male and female function (dichogamy) to promote autogamy, as well as reductions in traits associated with pollinator attraction and reward (Sicard and Lenhard 2011; Tsuchimatsu and Fujii 2022). For Lamiales families with TSSC, its loss appears to be a key additional component of this syndrome. TSSC has been lost in at least three (likely more) independent transitions to autogamous selfing within *Mimulus* (Friedman et al. 2017), as well as in selfing *Mazus pumilus* (Jin et al. 2015), consistent with rapid stigma closure functioning specifically in pollinator-mediated outcross male and/or female fitness.

We found a multi-genic basis for loss of stigma reactivity in selfer *M. nasutus,* with four significant QTLs together explaining ∼3/4 of the parental difference (Table 1). Each locus showed dominance toward one parent (Table 1), but there was little evidence of epistasis (Fig. 3). We take these estimates with a grain of salt, as semi-quantitative scoring of stigma closure violates some assumptions of QTL mapping models. However, a complex and cumulative genetic architecture for closure variation accords well with other observations as well (Burck 1902; Newcombe 1922). We only touched each stigma once for QTL mapping scores, but some slow or partially closed stigmas in hybrids will close fully with a second stimulation. Conversely, the stigmas of even very reactive lines become less sensitive and/or rapidly closing as individual flowers age (Milet-Pinheiro et al. 2009). Along with abundant variation in closure speed among the IM lines and among *Mimulus* populations (Meinke 1992; Friedman et al. 2017), this individual variability defines stigma closure as a truly quantitative trait. The two moderate QTLs for stigma closure detected in *M. parishii* x *M. cardinalis* hybrids (Chen et al., unpubl. MS), which do not as fully explain parental divergence and are in non-syntenic genomic regions, also suggest that TSSC is a complex quantitative trait presenting many targets for gradual loss. Together, these studies define a diverse but manageable core set of candidate genomic regions for further genetic dissection.

Mating system proxies and closure speed are correlated across multiple monkeyflower systems (Meinke 1992; Friedman et al. 2017), indicating that stigma reactivity decreases in concert with other traits defining the selfing syndrome. However, our results indicate that cross-population correlations are not due to a shared genetic basis (pleiotropy) or tight linkage in super-genes (Schwander et al. 2014). Except for *sc10*’s overlap with corolla width and length QTLs (*cw10, ct10*) near the *inv10* inversion and co-incidence of *sc6* with the *ff6* flowering time QTL, there is no evidence that the same genomic regions coordinate joint evolution of closure and other floral traits. Although *inv10* contributes to the cross-trait correlations in F_2_ hybrids, it is not a selfer supergene; it distinguishes Iron Mountain annual *M. guttatus* from most other *M. guttatus* accessions as well as *M. nasutus* (Flagel et al. 2019). Thus, QTL co-incidence there is likely a byproduct of suppression of recombination (in heterozygotes) among multiple genes with individually minor effects. Stigma closure was not associated with loci causing partial hybrid male sterility in the F_2_s (Table 1, Fig. 2). Beyond shared QTLs for corolla size metrics (Fishman et al. 2002) and pleiotropic side effects of hybrid anther sterility (Barr and Fishman 2011; Fishman et al. 2015), the selfing syndrome generally shows minimal genetic co-ordination in monkeyflower hybrids. In contrast, supergenes and/or structural variants are associated with life history strategies (Lowry and Willis 2010; Twyford and Friedman 2015), pollinator syndromes (Fishman et al. 2013; Liang et al. 2023), and edaphic adaptation (Toll and Willis 2023). A polygenic and un-coordinated genetic architecture for the selfing syndrome, extended here to stigma closure, likely reflects inbreeding’s promotion of linkage disequilibrium, abundant standing variation for mating system traits complex-wide, and a relatively flat and/or moving adaptive landscape for loss of pollinator-associated traits (e.g., Fishman et al. 2002, 2015).

Loss of TSSC could result from mutational accumulation under relaxed selection for its maintenance in selfers or (non-exclusively) from active selection for non-functionalization. In a prior selfer such as *M. nasutus* (Fig. 1c), stigmas risk self-triggering through physical contact with the anthers or corolla tube in bud. Reopening after insufficient pollination provides touch-sensitive stigmas with a second chance to increase pollen loads (Burck 1902; Newcombe 1922, 1924; Fetscher and Kohn 1999; Jin et al. 2015; Friedman et al. 2017). However, selfers may have highest fitness if they remain continuously open to self-pollination until all ovules are fertilized rather closing prematurely or attempting to actively “count” pollen tubes after closure. Thus, loss of both rapid TSSC and slower post-pollination closure (Fig. 1) may be actively favored in fully autogamous taxa to maximize seedset. Distinguishing neutral degeneration by drift from directional selection for loss is difficult, however, as mutations may be generally biased toward breakage (Tsuchimatsu and Fujii 2022). In addition, the stochastic loss of variation associated with the shift to selfing compromises short-term signatures of selection (Busch et al. 2022), as well as patterns of molecular evolution. Nonetheless, the consistent directionality of QTL effects (selfer alleles = slower at 6 out of 6 QTLs across this study and Chen et al., unpubl. MS) is consistent with active selection for non-closure (Orr 1998).

### The mechanistic basis of rapid touch-sensitive stigma closure and its loss

Stigma closure has long been recognized as a controlled wilt, parallel to the shifts in cell turgor that modulate stomatal closing or drive the rapid nastic movements of *Mimosa* leaflets and *Dionaea* flytraps (Hedrich and Neher 2018). In the latter cases of mechanosensitive movement, action potentials (Hedrich and Kreuzer 2023) are transmitted from the stimulated sensory cells to a specialized movement organ (e.g. pulvinus). There, directional changes in turgor pressure cause predictable changes in curvature that initiate organ movement, often amplified by buckling effects (Skotheim and Mahadevan 2005; Dumais and Forterre 2012; Mano and Hasebe 2021). Touch-sensitive stigmas, which move only a few millimeters, may be small enough to not require either a specialized pulvinus structure or snap-buckling to generate their rapid movement (Skotheim and Mahadevan 2005). Indeed, cells around a gentle touch point can be seen to shrink locally without precipitating full closure (Newcombe 1922). However, closure is not entirely linear -- slow but fully closing stigmas often “get stuck” and then visibly accelerate after the midway point with or without additional stimulation, suggesting snap-buckling by key clusters of cells.

Stigma closure thus necessarily involves turgor control and cell shape components as well as mechanosensing and signal transduction. These components interact with environmental factors (e.g. temperature, humidity) and flower age (Miyoshi 1891; Burck 1902; Newcombe 1922) to vary closure speed quantitatively, and each may be vulnerable to separate disruption to contribute to loss. The mechanosensing and signaling components of stigma closure were measured as action potentials in an early study of *Incarvillea* (Sinyukin and Britikov 1967), which also implicated pollen respiratory activity as the signal of successful pollination that blocks slow reopening. Turgor loss as the key to movement is evidenced by the observation that stigma lobes of excised *M. cardinalis* styles rapidly close (without touch) when the chamber they are in is placed under vacuum (Supplemental Figure/Video S1) and by the relatively slow (10-40 minutes) reopening of *Mimulus* stigmas as turgor is restored. Beyond these direct observations, our expression data provide some of the first clues into how stigma closure works and how it can evolve to not work. First, stigma epidermal cells could lose the capacity to mechanosense and/or transduce signals (i.e., the stigma does not know that it was touched). Second, stigma cells could remain turgid despite reception of the touch signal (i.e., the stigma knows that it was touched, but does not react). Third, stigma cells could maintain a constant shape regardless of turgor (i.e. the stigma knows and reacts to touch but cannot move). We consider candidate loci relevant to each of these stages below.

Genome-wide, we identified no “smoking gun” differences in SF *M. nasutus* or IM709 *M. guttatus* noncloser vs. *M guttatus* IM767 closer gene expression implicating major disruption of a known mechanosensing ion channel gene (Kurusu et al. 2013; Basu and Haswell 2017). However, an MSL10 family gene on Chr1 (Migut.A00554) was among the top-10 highest expressed style genes in all three lines and was 2-fold reduced in rare noncloser IM709. This gene was at most moderately expressed in the other seven tissues used for annotation of new IM62 v3, IM767 v1 and SF *M. nasutus* v2.1 reference genomes (https://phytozome-next.jgi.doe.gov/) as well as in leaf and stamen RNA samples paired with the style collections (L. Fishman, F.R. Finseth; unpubl. data). MSL8 helps maintain pollen turgor via cell wall strengthening (Wang et al. 2021; Miller et al. 2022) and parallel female reproductive functions during pollen tube growth may be the primary or ancestral function of the highly style-expressed *Mimulus* MSL10 gene. However, Migut.A00554 was one a handful of genes with extremely high style expression but relatively low peptide abundance in an IM *M. guttatus* stylar proteome (Aagaard et al. 2013); because that study excluded stigma tissue, this mismatch may indicate this MSL10 is specifically localized in the stigma. Intriguingly, the MSL10 group gene FLYCATCHER1 is similarly highly expressed in the trigger cells of the specialized prey-capture leaves of Venus flytraps and sundews (Procko et al. 2021). Thus, while the QTL mapping rules out genetic changes at Migut.A00554 as the cause of the recent loss of closure in *M. nasutus,* it remains a strong candidate mechanosensor for stigma closure. *Mimulus* is amenable to stable genetic manipulation (Yuan 2019), so functional tests of its role(s) are feasible to illuminate both stigma closure and reproductive mechanosensing more broadly.

As a local, rapid, and directional wilt, stigma closure depends on water movement out of some stigma/style cells and into others; thus, significant over-representation of aquaporins among genes with higher expression in the styles of noncloser *M. nasutus* suggests turgor maintenance as a mechanism Aquaporins allow for rapid bidirectional movement of water (and other solutes) across cell membranes, and thus are key components of anther dehiscence, pollen hydration, and pollen tube growth, as well as turgor maintenance in the face of abiotic stresses (Tyerman et al. 2021). Conceivably, increased density of aquaporins in the tonoplast and/or plasma membrane of noncloser styles may equalize turgor among stigma cells, preventing the sharp changes in pressure that lead to nastic movement. Intriguingly, calcium-dependent phosphorylation of a PIP (plasma intrinsic protein) aquaporin was recently implicated in the relatively slow turgor-driven opening and closing movements of gentian corollas in response to environmental cues (Dai et al. 2021; Nemoto et al. 2022). Thus, aquaporins may be key targets for further investigation of the cellular mechanisms of the loss of rapid stigma closure. Pharmacological inhibition of aquaporins (Tyerman et al. 2021), along with monitoring of turgor in individual cells across the genetic and environmental spectrum of *Mimulus* stigma closure (Beauzamy et al. 2014), allow functional validation of these hypotheses. By combining such approaches with finer mapping of turgor-related candidates within QTLs, we can assess whether genome-wide changes in aquaporin regulation are directly causal of closure loss or a downstream consequence of genetic disruption of other components.

Like stomatal guard cells (Kollist et al. 2014) or pulvini cells, stigma epidermal cells must shrink and re-swell unevenly to translate changes in turgor to directional movement. Thus, remodeling of cell walls to remove structural asymmetry or flexibility may be a simple step toward the secondary loss of rapid closure in selfer taxa. Patterns of gene expression may indicate cell wall stiffening in noncloser stigmas; in particular, the set of genes with higher expression in *M. nasutus* was enriched for aspartic protease/xylanase inhibitor functions, which regulate pectin deposition in expanding cell walls (Gao et al. 2016) and the set down in IM709 was enriched for xyloglucan endotransglycosylase/glycoside hydrolase family genes involved in cell wall degradation. In addition, a pectin methylesterase related to QUARTET 1 and a xyloglucan endotransglucosylase/hydrolase (XTH) were among the most DE in the interspecific contrast, with high expression in IM767 *M. guttatus* but essentially no reads in *M. nasutus.* QUARTET is essential for cell-wall softening to allow separation of pollen tetrads (Francis et al. 2006) and similar pectin modification enzymes reshape stomatal guard cells under heat stress (Amsbury et al. 2016). XTH enzymes modify cell-wall hemicelluloses and several touch-sensitive (TCH) genes in this family have been implicated in touch-induced changes in plant morphology (Braam 2005). Although neither of these genes is within a QTL, parallel (7-to 9-fold lower) expression shifts in the IM709 noncloser suggest that cell wall remodeling genes may mechanistically contribute to both losses of touch-sensitive movement.

### Conclusions

Overall, the genetic and transcriptomic analyses of the loss of rapid stigma closure suggest that this novel and complex plant movement trait involves sensing, turgor, and structural components built on deeper reproductive signaling systems in angiosperm styles. Repeated losses in diverse selfers, independent genomic bases for loss in *M. nasutus* and *M. parishii* (Chen et al., unpubl. MS), and a well-resolved multi-locus architecture in *M. nasutus* x *M. guttatus* hybrids provide multiple evolutionary and functional paths to understanding stigma closure as both a key component of the selfing syndrome and as a novel Lamiales trait. Parallel shifts in style expression patterns in selfer *M. nasutus* and rare *M. guttatus* noncloser IM709 suggest introgression and/or shared downstream targets during loss of TSSC, as well as revealing other reproductive trait variation (e.g. changes in volatiles associated with defense and/or pollinator attraction). Resources in monkeyflowers, including stable transformation protocols, multiple new reference genomes annotated with transcriptomes from diverse tissues, and reproductive proteomes, make *Mimulus* an ideal model for further mechanistic investigation. Thus, our findings provide a strong foundation for reconstruction of the origins and mechanisms of an evolutionarily novel plant behavior, as well as new insight into plant reproductive diversification.

## Author contributions

Mariah McIntosh and Kailey Baesen; investigation, editing. Thomas Nelson; formal analysis, methodology, drafting. Findley R. Finseth; conceptualization, editing, resources, funding acquisition. Evan Stark-Dykema; analysis, data curation, drafting, editing. Lila Fishman; conceptualization, supervision, investigation, analysis, drafting and editing, visualization, and funding acquisition.

## Acknowledgments

We thank David Denghui Xing of the UM Genomics Core and Andrew Demaree for assistance with laboratory projects and October Moynahan of the UM ECOR Plant Growth Facility, Colette Berg, and Natalie Dietz for assistance with plant care. We are grateful to Timothy Wheeler for his imaging of flowers, to Brandon Cooper for sharing his camera set-up and to Joshua Puzey for allowing use of his videos of stigma closure.

## Data accessibility statement

New genomic and transcriptomic datasets presented in this paper are archived on the NCBI Sequence Archive as projects PRJNA1063293 and PRJNA1063748, respectively. The mapping population genotypes and phenotypes, as well as RNASeq readcounts and related data, are archived on Dryad at doi:10.5061/dryad.mw6m9063v.

