## Supplemental File 1 - Table S1 and Video S1 legend for "Undoing the ‘nasty: dissecting touch-sensitive stigma movement (thigmonasty) and its loss in self-pollinating monkeyflowers"

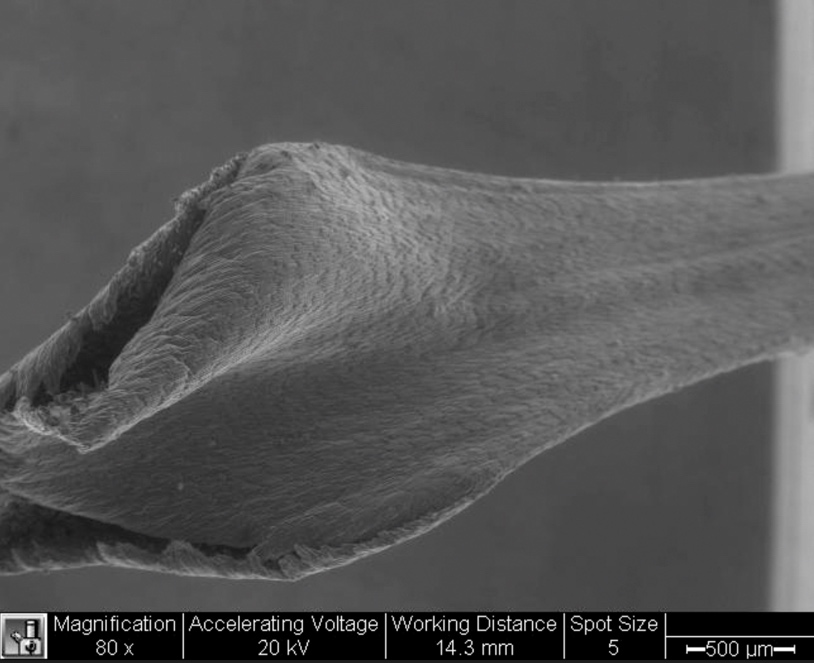

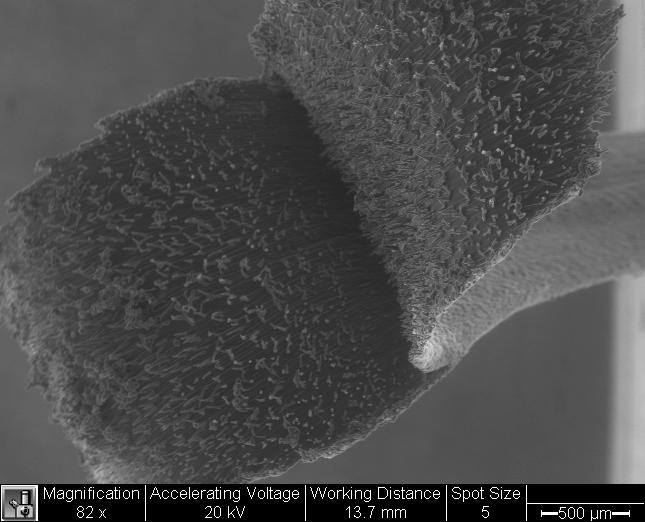
**Figure/Video S1.** A. Scanning electron microscopy (SEM) video (double-click to run) of an excised monkeyflower style showing stigma closure under vacuum. B. Still image of the closed stigma after 2 seconds. Video credit: Joshua Puzey. The video is also supplied as Supplemental Video 1 in .mov format.

**Table S1.** Genes with top-50 ranked expression in stylar (including stigma) tissue of at least one of the three yellow monkeyflower genotypes. Genes are ordered by rank in the IM767 *M. guttatus* line with fast stigma closure and annotated with their *Arabidopsis thaliana* best hit gene and its description (defline). Where a gene was identified as present in the style (not including stigma) proteome of IM62 *M. guttatus* (Aagaard et al. 2013), the relative peptide abundance (MuDPIT NSAF value) is provided for comparison. Dashes indicate a v2 gene was not identified as present in the style proteome, but a *de novo* analysis of peptides aligned to newer genome assemblies/annotations was not conducted.

| ***M. guttatus***  **v2 gene name** | ***SF M. nasutus* readcount^a^** | **IM709**  ***M. guttatus* readcount^a^** | **IM767**  ***M. guttatus* readcount^a^** | ***Arabidopsis***  **best hit** | ***Arabidopsis* defline** | **Proteome**  **MuDPIT NSAF** |
| --- | --- | --- | --- | --- | --- | --- |
| Migut.M01363 | 82900 | 49742 | 85906 | AT1G05010 | ethylene-forming enzyme | 0.001046 |
| Migut.J00046 | 69184 | 36278 | 65709 | AT3G51590 | lipid transfer protein 12 | 0.000903 |
| Migut.E01125 | 14467 | 21098 | 65543 | AT4G11650 | osmotin 34 | 0.000640 |
| Migut.C00884 | 77118 | 41079 | 64152 | AT1G20160 | Subtilisin-like serine endopeptidase family protein | 0.019309 |
| Migut.F00699 | 68826 | 35489 | 53915 | AT4G00430 | plasma membrane intrinsic protein 1;4 | - |
| Migut.G00720 | 1789 | 466 | 38226 | AT5G23960 | terpene synthase 21 | 0.000635 |
| Migut.A00554 | 38635 | 18440 | 37750 | AT5G12080 | mechanosensitive channel of small conductance-like 10 | 0.000076 |
| Migut.N01583 | 24446 | 54047 | 33973 | AT2G45600 | alpha/beta-Hydrolases superfamily protein | 0.020304 |
| Migut.J00043 | 20639 | 12355 | 33805 | AT5G59310 | lipid transfer protein 4 | 0.000339 |
| Migut.F00292 | 34329 | 40778 | 31436 | AT3G48950 | Pectin lyase-like superfamily protein | 0.002513 |
| Migut.H00102 | 15426 | 23242 | 27774 | AT3G27810 | myb domain protein 21 | - |
| Migut.J00549 | 56886 | 843 | 27145 | AT3G53420 | plasma membrane intrinsic protein 2A | 0.000985 |
| Migut.F01647 | 7756 | 11659 | 26608 | AT5G16010 | 3-oxo-5-alpha-steroid 4-dehydrogenase family protein | 0.000146 |
| Migut.E00478 | 21811 | 24899 | 25074 | AT2G02100 | low-molecular-weight cysteine-rich 69 | - |
| Migut.M01454 | 17966 | 17739 | 24676 | AT4G39090 | Papain family cysteine protease | 0.000500 |
| Migut.O00218 | 22267 | 12732 | 23879 | AT5G63180 | Pectin lyase-like superfamily protein | 0.000098 |
| Migut.D00373 | 12498 | 15132 | 23662 | AT4G14550 | indole-3-acetic acid inducible 14 | - |
| Migut.J00330 | 29461 | 23858 | 21486 | AT3G09390 | metallothionein 2A | - |
| Migut.G00850 | 10612 | 10136 | 20073 | AT1G05680 | Uridine diphosphate glycosyltransferase 74E2 | 0.000499 |
| Migut.D02284 | 27248 | 15293 | 19398 | AT3G45010 | serine carboxypeptidase-like 48 | 0.000494 |
| Migut.B01762 | 30228 | 16484 | 19138 | AT2G45960 | plasma membrane intrinsic protein 1B | - |
| Migut.N01325 | 25370 | 10281 | 18821 | AT4G35100 | plasma membrane intrinsic protein 3 | 0.001700 |
| Migut.H00923 | 7440 | 16874 | 18353 | AT5G15230 | GAST1 protein homolog 4 | - |
| Migut.D01550 | 5875 | 808 | 18228 | AT5G23960 | terpene synthase 21 | 0.000130 |
| Migut.H01595 | 30780 | 31240 | 18183 | AT1G05010 | ethylene-forming enzyme | 0.001044 |
| Migut.C00509 | 13602 | 20801 | 17975 | AT5G59845 | Gibberellin-regulated family protein | - |
| Migut.A00823 | 21015 | 23855 | 17649 | AT4G12500 | Bifunctional inhibitor/lipid-transfer protein | 0.000730 |
| Migut.C00231 | 5710 | 10886 | 16658 | AT2G30860 | glutathione S-transferase PHI 9 | 0.000186 |
| Migut.H01994 | 56740 | 23821 | 16419 | AT5G59310 | lipid transfer protein 4 | - |
| Migut.F00332 | 43476 | 29618 | 16254 | AT5G20390 | Glycosyl hydrolase superfamily protein | 0.011539 |
| Migut.B00929 | 14137 | 21860 | 15989 | AT3G59850 | Pectin lyase-like superfamily protein | - |
| Migut.C00923 | 24738 | 7036 | 14732 | AT1G76180 | Dehydrin family protein | - |
| Migut.G00723 | 1004 | 1526 | 14631 | AT5G23960 | terpene synthase 21 | - |
| Migut.J00768 | 11209 | 5036 | 14600 |  |  | - |
| Migut.H02487 | 13110 | 9952 | 14408 | AT2G33170 | Leucine-rich repeat receptor-like protein kinase family protein | 0.000188 |
| Migut.D01172 | 2283 | 2716 | 14169 | AT5G61820 |  | - |
| Migut.J01425 | 1164 | 13588 | 13999 | AT4G30380 | Barwin-related endoglucanase | 0.000936 |
| Migut.N00045 | 4986 | 2016 | 13660 | AT3G36659 | Plant invertase/pectin methylesterase inhibitor superfamily | - |
| Migut.K00277 | 6370 | 9497 | 13449 | AT5G57800 | Fatty acid hydroxylase superfamily | - |
| Migut.C00776 | 6017 | 8266 | 13122 | AT3G51030 | thioredoxin H-type 1 | 0.000083 |
| Migut.K00920 | 4085 | 7473 | 13011 | AT5G37600 | glutamine synthase clone R1 | 0.000900 |
| Migut.F01419 | 30381 | 13283 | 12951 | AT4G00430 | plasma membrane intrinsic protein 1;4 | 0.000884 |
| Migut.C01407 | 19735 | 3786 | 12846 | AT1G33430 | Galactosyltransferase family protein | 0.000398 |
| Migut.M01402 | 22796 | 11918 | 12596 |  |  | 0.006074 |
| Migut.A00666 | 4479 | 6587 | 12584 |  |  | - |
| Migut.L00106 | 7395 | 4365 | 12576 | AT4G08685 | Pollen Ole e 1 allergen and extensin family protein | - |
| Migut.K00035 | 3624 | 7914 | 12375 | AT2G37620 | actin 1 | 0.006747 |
| Migut.I00379 | 625 | 4713 | 12049 | AT5G26330 | Cupredoxin superfamily protein | - |
| Migut.N02028 | 6523 | 19002 | 11910 | AT3G51895 | sulfate transporter 3;1 | 0.000061 |
| Migut.J00325 | 8317 | 18322 | 11591 | AT2G39700 | expansin A4 | 0.001118 |
| Migut.D00153 | 17692 | 23381 | 11452 | AT5G23660 | homolog of Medicago truncatula MTN3 | - |
| Migut.F00138 | 15588 | 15297 | 11237 | AT5G24580 | Heavy metal transport/detoxification superfamily protein | - |
| Migut.L01053 | 16452 | 8980 | 11060 | AT1G25330 | basic helix-loop-helix (bHLH) DNA-binding superfamily | - |
| Migut.J00202 | 13645 | 4812 | 10856 | AT5G01210 | HXXXD-type acyl-transferase family protein | 0.000421 |
| Migut.H01853 | 10246 | 11219 | 10721 | AT4G05320 | polyubiquitin 10 | - |
| Migut.F02008 | 27645 | 20721 | 10489 | AT3G54820 | plasma membrane intrinsic protein 2;5 | 0.001043 |
| Migut.N03349 | 5090 | 10778 | 10460 | AT1G71950 | Proteinase inhibitor, propeptide | 0.002792 |
| Migut.B00928 | 3441 | 18791 | 10190 | AT2G43870 | Pectin lyase-like superfamily protein | - |
| Migut.H01669 | 8458 | 13971 | 9984 | AT1G79550 | phosphoglycerate kinase | 0.001273 |
| Migut.J00047 | 5494 | 11570 | 9454 | AT3G08770 | lipid transfer protein 6 | - |
| Migut.D02291 | 21947 | 8062 | 9214 | AT2G36830 | gamma tonoplast intrinsic protein | - |
| Migut.L01795 | 14164 | 6402 | 9143 | AT3G52610 |  | - |
| Migut.I00928 | 16652 | 14626 | 9020 | AT3G16240 | delta tonoplast integral protein | 0.000179 |
| Migut.L01923 | 15898 | 2629 | 8375 | AT5G36110 | cytochrome P450, family 716, subfamily A, polypeptide 1 | 0.000028 |
| Migut.G01285 | 992 | 15653 | 8302 | AT3G51590 | lipid transfer protein 12 | 0.001919 |
| Migut.N01231 | 2954 | 14372 | 8177 | AT1G49310 |  | - |
| Migut.D01302 | 19098 | 4026 | 6945 | AT4G25040 | Uncharacterized protein family (UPF0497) | - |
| Migut.N00127 | 5355 | 11107 | 4569 | AT1G08830 | copper/zinc superoxide dismutase 1 | - |
| Migut.B00257 | 28236 | 2407 | 4195 | AT3G18830 | polyol/monosaccharide transporter 5 | 0.000080 |
| Migut.A00738 | 20108 | 11690 | 3627 | AT4G00430 | plasma membrane intrinsic protein 1;4 | 0.001010 |
| Migut.A00821 | 2085 | 29270 | 3605 | AT4G12520 | Bifunctional inhibitor/lipid-transfer protein | - |
| Migut.N00668 | 3591 | 10691 | 2726 | AT4G12910 | serine carboxypeptidase-like 20 | 0.003382 |
| Migut.M00643 | 1692 | 11680 | 2720 |  |  | 0.005844 |
| Migut.N02885 | 3968 | 11507 | 2585 | AT4G33510 | 3-deoxy-d-arabino-heptulosonate 7-phosphate synthase | 0.000568 |
| Migut.J00813 | 14247 | 6887 | 2278 | AT2G25630 | beta glucosidase 14 | 0.001965 |
| Migut.D02133 | 17925 | 1637 | 2043 | AT4G01470 | tonoplast intrinsic protein 1;3 | - |
| Migut.G01288 | 16222 | 2687 | 2005 | AT5G59310 | lipid transfer protein 4 | - |
| Migut.N01232 | 29312 | 4707 | 1990 |  |  | - |
| Migut.A00217 | 14255 | 4024 | 1647 | AT2G25810 | tonoplast intrinsic protein 4;1 | 0.000162 |
| Migut.J00672 | 13392 | 2015 | 1503 | AT2G47140 | NAD(P)-binding Rossmann-fold superfamily protein | - |
| Migut.J00462 | 40902 | 585 | 1207 | AT1G75800 | Pathogenesis-related thaumatin superfamily protein | - |
| Migut.M01405 | 1214 | 11145 | 313 | AT5G47560 | tonoplast dicarboxylate transporter | - |
| Migut.J00756 | 102948 | 16153 | 306 | AT5G59320 | lipid transfer protein 3 | - |
| Migut.F01480 | 13843 | 1668 | 241 | AT4G37430 | cytochrome P450, family 91, subfamily A, polypeptide 2 | - |
| Migut.M00388 | 21076 | 24 | 34 | AT4G15210 | beta-amylase 5, ram1 | - |

^a^Readcounts are means of counts in the batch-normalized replicates (n = 2 for SF, 3 for others) of each genotype, standardized by gene length.
